# Empirical Bayes estimation of semi-parametric hierarchical mixture models for unbiased characterization of polygenic disease architectures

**DOI:** 10.1101/080945

**Authors:** Jo Nishino, Yuta Kochi, Daichi Shigemizu, Mamoru Kato, Katsunori Ikari, Hidenori Ochi, Hisashi Noma, Kota Matsui, Takashi Morizono, Keith A Boroevich, Tatsuhiko Tsunoda, Shigeyuki Matsui

## Abstract

Genome-wide association studies (GWAS) suggest that the genetic architecture of complex diseases consists of unexpectedly numerous variants with small effect sizes. However, the polygenic architectures of many diseases have not been well characterized due to lack of simple and fast methods for unbiased estimation of the underlying proportion of disease-associated variants and their effect-size distribution. Applying empirical Bayes estimation of semi-parametric hierarchical mixture models to GWAS summary statistics, we confirmed that schizophrenia was extremely polygenic (∼ 40% risk variants of independent genome-wide SNPs, most within odds ratio (OR)=1.03), whereas rheumatoid arthritis was less polygenic (∼ 4 to 8% risk variants, significant portion reaching OR=1.05 to 1.1). For rheumatoid arthritis, stratified estimations revealed that expression quantitative loci in blood explained large genetic variance, and low- and high-frequency derived alleles were prone to be risk and protective, respectively, suggesting a predominance of deleterious-risk and advantageous-protective mutation. Despite genetic correlation, effect-size distributions for schizophrenia and bipolar disorder differed across allele frequency. These analyses distinguished disease polygenic architectures and provided clues for etiological differences in complex diseases.

---

Genome-wide association studies (GWAS) have identified numerous susceptibility variants for complex diseases1. The sets of variants identified from GWAS, however, can generally explain only a small proportion of the heritability estimated from family studies, the so called “missing heritability” problem2. Many researches have suggested that the variance explained by all SNPs in dense genotyping arrays, i.e., SNP heritability, often accounts for a large proportion of the family-based heritability^3–11^.

Quantitative evaluation of the polygenic architecture, in particular, the estimation of the proportion of disease-associated SNPs and their effect-size distribution, is essential to further determine the source of observed heritability^7,8,12–16^. The estimation of these components also contributes to accurate power and sample size calculations of GWAS^7,12,13,17–19^ and estimation of the predictive capability of disease risks^12,15,16^.

However, we are still far from understanding the polygenic architecture of most complex diseases, because so far, there have been no feasible or fast methods to unbiasedly evaluate various polygenic architectures using the entire SNPs across the genome. Stahl et al. proposed estimating the proportion of disease-associated SNPs and the effect-size distribution using an approximate Bayesian polygenic analysis^8^. Its application, however, has been limited to few studies^7,8^ because of technical complexity and excess computational burden with many simulations. On the other hand, some authors estimated the effect-size distribution based on a power evaluation for SNPs reaching genome-wide significance^13–15^. This method, however, is to evaluate effect sizes only for those SNPs with relatively large effects, not all the disease-associated SNPs, requiring adjustment for the winner’s curse (selection bias in using top significant SNPs) in the effect-size estimation.

To address the aforementioned limitations of the existing methods, we propose an empirical Bayes estimation of semi-parametric hierarchical mixture models (SP-HMMs)^20,21^ of GWAS summary statistics on effect sizes, such as estimated log-odds ratios to associate genotypes with disease susceptibility (Online Methods). Mixture modelling refers to decomposing the underlying distribution of SNP-specific summary statistics into a non-null distribution for SNPs associated with disease occurrence, which corresponds to a signal component, and a null distribution for the remaining SNPs without association, which corresponds to a noise component, with a mixing probability or proportion of disease-associated SNPs, *π*. For non-null distribution, semi-parametric hierarchical modelling incorporates standard asymptotic normality for summary statistics, while the true effect sizes follow a non-parametric prior distribution *g*. With an expectation-maximization (EM) algorithm^22^, we can unbiasedly estimate the prior probability *π* and distribution *g* using the data, i.e., empirical Bayes estimation. The empirical Bayes estimation of hierarchical mixture models is also applicable for SNP heritability estimation^3^ and adjustment for the winner’s curse^23^.

The features of our approach are summarized as follows: 1) the polygenic architecture for the entire set of SNPs, represented by *π* and *g*, can be flexibly and unbiasedly estimated, 2) it requires only summary data from GWAS (e.g., estimated log-odds ratios and standard error for individual SNPs are used), and 3) the estimation algorithm is easily implemented and fast.

Throughout this paper, we fit the SP-HMM to summary data from meta-/mega-analyses for various diseases such as rheumatoid arthritis^24^, schizophrenia^25^, bipolar disorder^26^, and coronary artery disease^27,28^, to estimate the respective polygenic architectures and compare them across diseases. We also assess the liability-scale variance explained by SNPs based on this estimation. In order to obtain further insight into the underlying polygenic architectures, our approach can be applied to SNPs belonging to important functional categories, such as expression quantitative trait loci (eQTL), coding, non-synonymous, promoter, 5’ or 3’ UTR, enhancer, and DNase I hypersensitivity sites^29–32^. We focus on eQTLs, as gene expression levels have been increasingly recognized as notable endophenotypes or important mediators between genetic variations and disease phenotypes^29,30,33,34^. Lastly, we also applied our method to GWAS data stratifying by derived allele frequency (DAF), rather than minor allele frequency (MAF)14,35,36. A minor allele with low MAF can represent an allele with high DAF possibly under positive selection, as well as an allele that is maintained at low DAF by negative selection. Thus, our DAF-based analysis facilitates interpretation from the perspective of population genetics^37^, possibly contributing to further understanding of the genetic etiology for complex diseases.

## RESULTS

We first confirmed the adequacy of our estimation method in unbiasedly estimating the proportion of disease-associated SNPs, *π*, and their effect-size distribution, *g*, in simulation experiments (Supplementary Note; Supplementary Table 1-2; Supplementary Fig. 1-17).The non-parametric estimation for *g* could flexibly capture various forms of the underlying effect-size distributions.

**Table 1.**
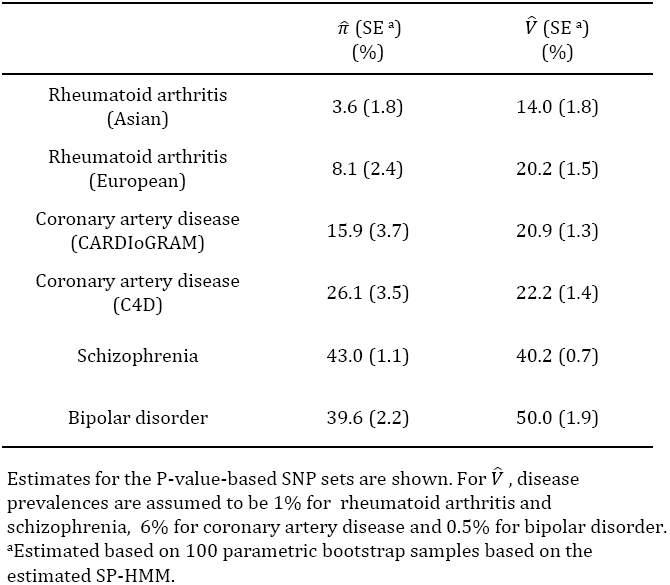
Estimated proportions of disease-associated SNPs, 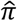, and liability-scale variance explained by SNPs, 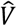

| | $\hat{\pi}$ (SE <sup>a</sup>)<br>(%) | $\hat{V}$ (SE <sup>a</sup>)<br>(%) |
| --- | --- | --- |
| Rheumatoid arthritis<br>(Asian) | 3.6 (1.8) | 14.0 (1.8) |
| Rheumatoid arthritis<br>(European) | 8.1 (2.4) | 20.2 (1.5) |
| Coronary artery disease<br>(CARDIoGRAM) | 15.9 (3.7) | 20.9 (1.3) |
| Coronary artery disease<br>(C4D) | 26.1 (3.5) | 22.2 (1.4) |
| Schizophrenia | 43.0 (1.1) | 40.2 (0.7) |
| Bipolar disorder | 39.6 (2.2) | 50.0 (1.9) |

Estimates for the P-value-based SNP sets are shown. For $\hat{V}$ , disease prevalences are assumed to be 1% for rheumatoid arthritis and schizophrenia, 6% for coronary artery disease and 0.5% for bipolar disorder.
<sup>a</sup>Estimated based on 100 parametric bootstrap samples based on the estimated SP-HMM.

**Figure 1.**
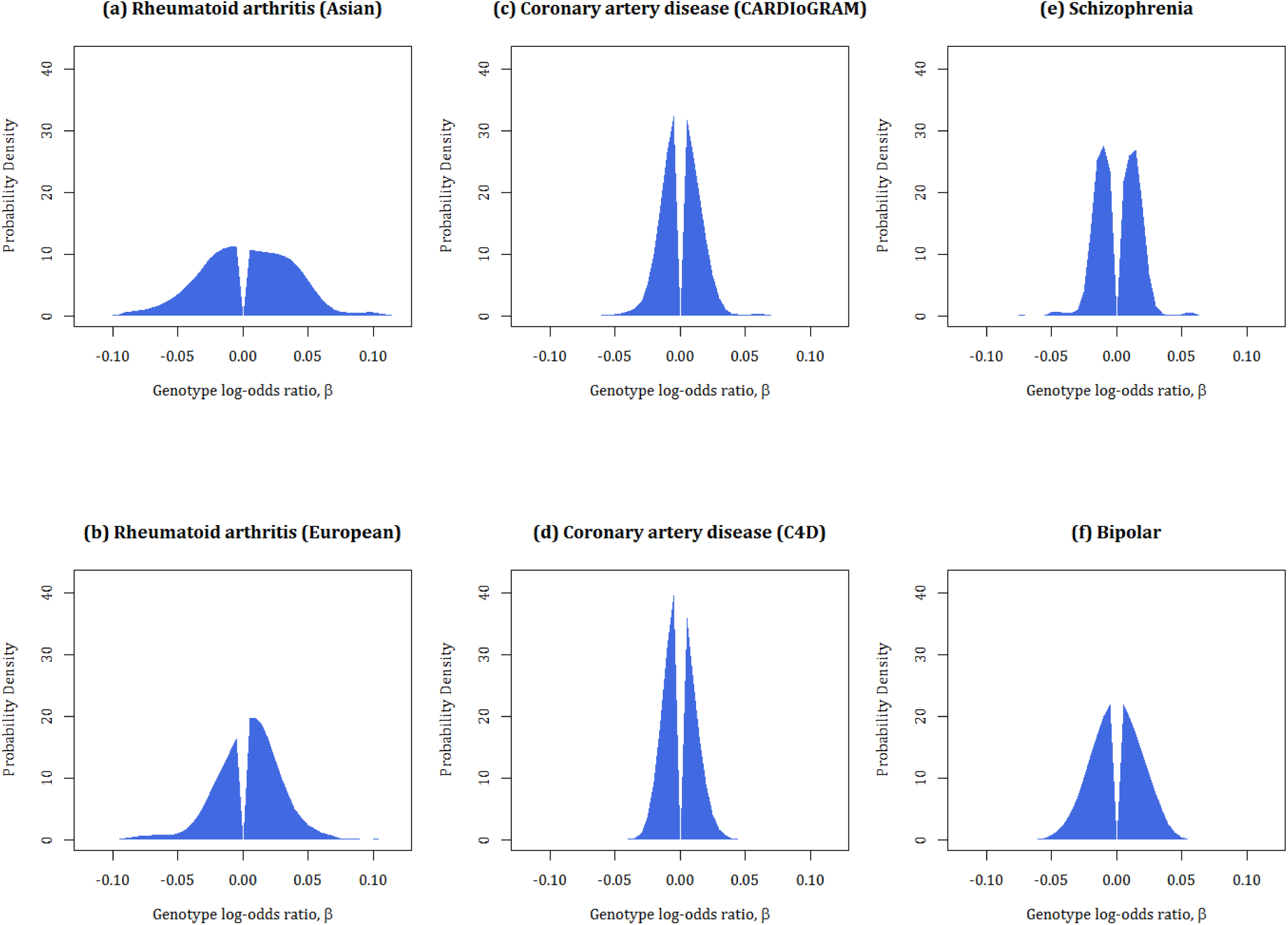
Estimated effect-size distributions for disease-associated SNPs, 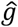. The P-value-based pruned SNP sets are used.

For application to real GWAS datasets, we used publicly available summary statistics from large meta-/mega-GWAS for the four complex diseases (see Supplementary Tables 3 and 4 for details of the GWAS data). In associating each genotype with disease susceptibility, we defined the effect size as a log-odds ratio of the derived allele relative to the ancestral allele, denoted by *β*. We obtained an estimate of *β* and its variance estimate from the summary data. The ancestral/derived alleles for each SNP were determined from dbSNP.

### Estimated proportion of disease-associated SNPs and effect-size distribution

To estimate the proportion of disease-associated SNPs, *π*, and the effect-size distribution, *g*, based on independent SNPs, we used two pruned SNP sets: P-value-based and random-pruned sets. The P-value-based method preferentially selected SNPs with stronger associations (hence more closely linked to causal variants) while using other GWAS data to correct for selection bias (see Online Methods for details). The random-pruned method sampled SNPs randomly. In both methods, pruned SNPs with linkage disequilibrium (LD, *r*^2^ ≤0.1) were selected. Of note, one causal variant would not be redundantly tagged by SNPs in the pruned SNP sets, whereas not all causal variants would be well tagged by SNPs even in the P-value-based sets. Thus, the estimates 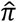 × (the number of SNPs in the SNP sets) using the pruned sets would give conservative estimates of the number of causal variants.

We fit the SP-HMM to the P-value-based pruned SNP sets in each GWAS (Table 1; Fig. 1). For rheumatoid arthritis, *π* was estimated as 3.6% for Asian and 8.1% for European populations, which were lower than the other diseases. The estimates of *π* were larger for two psychiatric diseases: 43.0% for schizophrenia and 39.6% for bipolar disorder. For coronary artery disease, using CARDIoGRAM and C4D data, *π* was estimated to be 15.9% and 26.1%, respectively.

With regard to the estimation of *g*, rheumatoid arthritis was shown to have a significant portion with larger effects, spanning to |*β*| = 0.05 (odds ratio of 0.95 or 1.05) or larger (Fig. 1). It is noteworthy that, for rheumatoid arthritis, the proportion of positive effects was clearly larger than that of negative effects, indicating that the derived alleles are more likely to be risk alleles for the disease. Bipolar disorder was also estimated to have a distribution with relatively large effects. In contrast, schizophrenia and coronary artery disease was shown to have narrower distribution with very small effects. Schizophrenia was shown to have peaks around |*β*| = 0.05.

The estimates of *π* for the random-pruned SNP sets were similar to those for the P-value-based SNP sets for each GWAS (Supplementary Table 5). For the estimation of effect-size distribution, 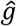 the absolute effect size, |*β*|, tended to be slightly greater when using the P-value-based SNP sets than when using the random-pruned SNP set (Supplementary Fig. 18).

### Liability-scale variance explained by SNPs

Using the estimates of the polygenic architecture (*π* and *g*), together with disease prevalence and allele frequencies, we could immediately evaluate the liability-scale variance, *V*, explained by SNPs (i.e., SNP heritability). For evaluating *V*, the SP-HMM could directly model binary traits (i.e., disease occurrence) via log-odds ratios obtained from GWAS summary data.

Using the P-value-based pruned SNP sets, for rheumatoid arthritis, the estimates of *V* were 14.0% for Asian and 20.2% for European data (Table 1). Based on the estimated variance of 12% explained by the major histocompatibility complex (MHC) region (removed from the SNP set) and family based heritability of 55% (Supplementary Table 1 of Stahl et al.8), SNPs explained 47.3% (= (0.14 + 0.12)/0.55) and 58.2% (= (0.20 + 0.12)/0.55) of the family based heritability for the Asian and European populations, respectively, which were generally consistent with the previous estimate of 65%8. The estimates of *V* in schizophrenia and bipolar disorder were 40.2% and 50%, respectively, which were higher but almost within the range of previously reported estimates of 23-43% and 37-47%, respectively, for these diseases^5,7,9,10^. For cardiovascular disease, the estimates of *V* from the CARDIoGRAM and C4D data were 20.9% and 22.2%, respectively.

The estimates of *V* for the P-value-based pruned SNP sets (Table 1) were greater than those for the random-pruned SNP sets, but the differences were not substantial except for bipolar disorder (Supplementary Table 5).

### Stratified estimation for eQTL/non-eQTL-SNPs

In order to gain insights into mediator effects of gene expression level, we fit the SP-HMM to ‘eQTL’ SNPs, detected as cis-eQTLs using peripheral blood samples^38^, and the remaining ‘non-eQTL’-SNPs, separately (Fig. 2). All the SNPs in this analysis were selected to be nearly independent using a LD-pruning method based on LD (*r*^2^ > 0.1) (see Online Methods).

**Figure 2.**
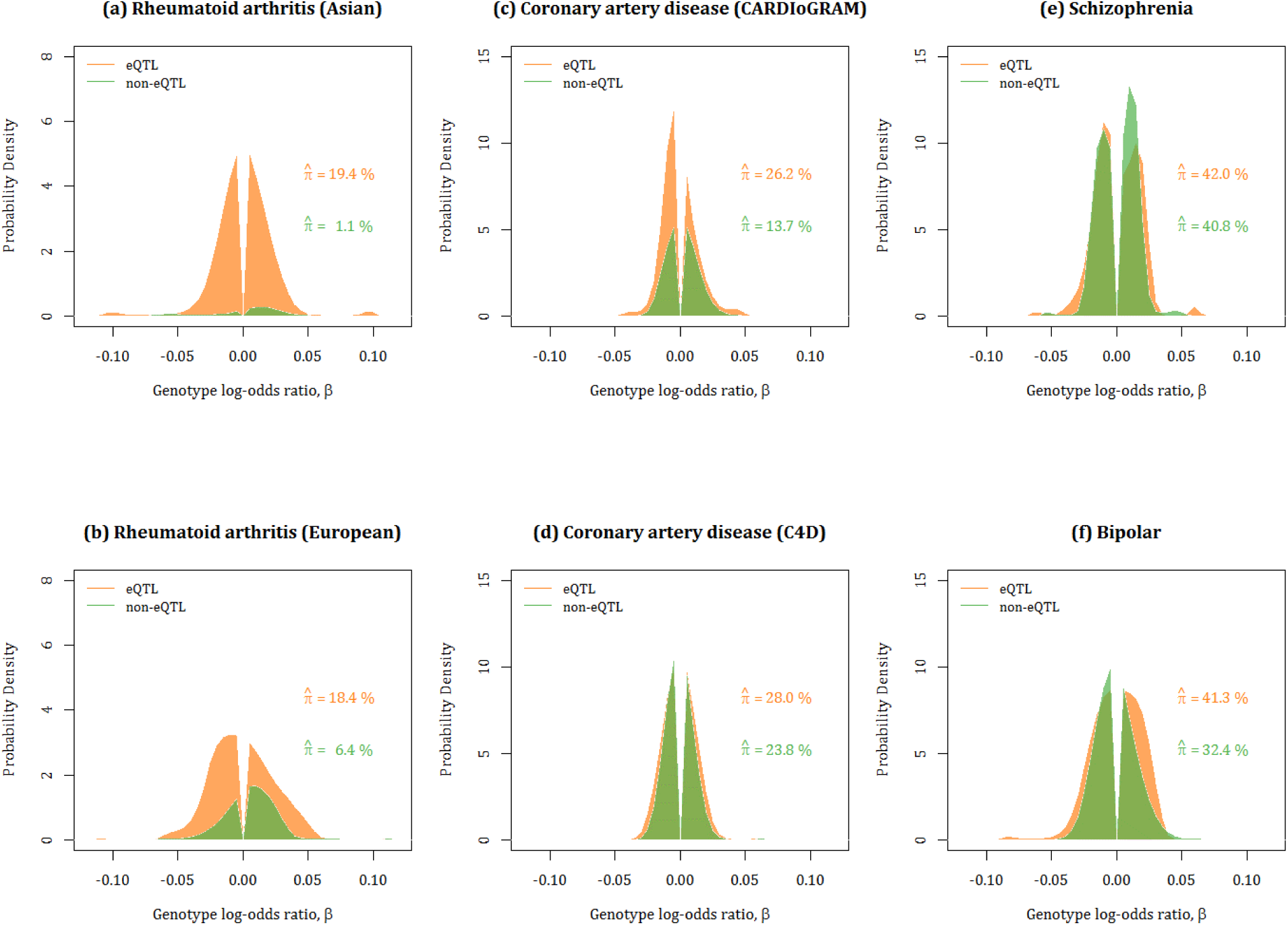
Estimated effect size distributions for eQTL-SNPs and non-eQTL-SNPs, 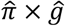. Green and orange lines show the results for the eQTL-SNP and non-eQTL-SNP sets, respectively. Estimated proportions of disease-associated SNPs, 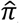, correspond to the areas under the curves.

For rheumatoid arthritis in Asian and European populations, the proportions of disease-associated SNPs in the eQTL-SNPs were estimated to be larger than that in the non-eQTL-SNPs (Fig. 2). In addition, the estimated effect-size distributions in terms of *π* × *g* (frequencies in the entire set including both null and non-null SNPs) in Fig. 2 indicated that there was a significant portion of SNPs with large effects, |*β*| > 0.05, for the eQTL-SNPs, but a small portion for the non-eQTL-SNPs, suggesting that the set of eQTL-SNPs included more components with distinctive large effects for rheumatoid arthritis. For the other diseases, there was a tendency for the frequencies of disease-associated SNPs in the set of eQTL-SNPs to be larger than those of the non-eQTL-SNPs.

We also estimated *V* for the eQTL-SNPs and non-eQTL-SNPs, separately (Supplementary Table 6). For rheumatoid arthritis, as expected from Fig. 2, the per-SNP variance for the eQTL-SNPs was much larger than for the non-eQTL-SNPs. Interestingly, although ‘eQTL’ was defined using European samples^38^, the enrichment of per-SNP variance (10.7-fold) in the eQTL-SNPs in the Asian population was larger than the 5.2-fold enrichment seen in the European population.

### Estimation across derived allele frequencies

The effect size estimation of GWAS data stratified with the derived allele frequency (DAF) could provide another perspective on polygenic architecture, which facilitates assessment based on population genetics (see Discussion). We classified all SNPs into five equally-sized DAF bins and estimated the effect-size distribution for each bin. For rheumatoid arthritis, the estimated distributions across the DAF bins were similar between Asian and European data (Figs. 3a, b). We observed peaks at positive effects, i.e., *β* > 0, for lower DAF bins, especially for DAF ≤0.2, and at negative effects for higher DAF bins, especially for DAF > 0.8. This indicates that low-frequency-derived and high-frequency-derived alleles are prone to act as risk and protective variants for disease occurrence, respectively. For coronary artery diseases, there was no substantial difference in the estimated effect-size distribution among DAF bins, compared with rheumatoid arthritis. For schizophrenia and bipolar disorder, we observed opposite tendencies: for schizophrenia, positive and negative effects were over-represented, especially at DAF < 0.2 and DAF > 0.8, respectively, whereas, for bipolar disorder, negative and positive effects were over-represented at DAF ≤0.2 and DAF > 0.8, respectively.

## DISCUSSION

We have developed a simple and fast method for unbiasedly estimating the proportion of disease-associated variants and the effect-size distribution based on the empirical Bayes estimation of SP-HMM. The proposed method can effectively distinguish various polygenic architectures, including the degree of polygenicity, across diseases, and can also provide various perspectives of the polygenic architecture based on important variant categories such as DAF and eQTL.

Schizophrenia, which has been suspected to be highly polygenic^39,40^, was estimated to have ∼ 40% disease-associated variants with very small effects (most within |*β*| = 0.03) of independent SNPs in the genome (Table 1, Fig. 1, Supplementary Table 5, and Supplementary Fig. 19). This suggests at least ∼ 40,000 causal variants exist in the genome, which does not contradict a recent study that estimated at least ∼ 20,000 causal variants by a simulation-based method^41^. The highly polygenicity of schizophrenia have been also confirmed by the observation that local SNP heritability estimates in independent LD blocks for schizophrenia were the most ubiquitously distributed among seven complex diseases^42^. In contrast, rheumatoid arthritis was found to be less polygenic. Our estimates of *π*, 3.6% for Asians and 8.1% for Europeans, were generally consistent with previous estimates of 2.7%8 or 5.4%10 for Europeans, and the significant portion of the estimate for *g* ranged to |*β*| = 0.05 or even 0.1 (Table 1 and Fig. 1). In fact, the effect sizes of validated variants for rheumatoid arthritis were generally larger than those for schizophrenia^24,25^. Our estimate of *g* means that the effect sizes of variants that will be detected in future would also be relatively larger among complex diseases. For coronary artery disease, the degree of polygenicity was estimated to be between those of rheumatoid arthritis and the two psychiatric diseases. The estimate of *π* for CARDIoGRAM was larger than that for C4D particularly in the P-value based pruned SNP set. Since SNPs of C4D were pruned by using LD structure of European ancestry (see Online Method), LD remaining in Asian SNPs, possibly linked with one causal variant, might put upward the proportion of disease-associated SNP.

In the rheumatoid arthritis stratification analysis based on eQTLs, we observed a high enrichment of per-SNP variance due to eQTLs determined by peripheral blood samples (Supplementary Table 6), similar to the enrichment on per-SNP variance by blood-specific DNaseI hypersensitivity sites (DHS)33, which were also strongly associated with expression variation^43^. As peripheral blood samples include multiple types of leukocytes, the eQTLs have the potential to control immune-related gene expressions that are associated with the occurrence of rheumatoid arthritis. Although ‘eQTL’ was defined using European samples^38^, the enrichment of 10.7-fold in the Asian population was larger than the 5.7-fold enrichment observed in the European population. The same tendency has been observed for the validated 100 non-MHC SNPs (Extended Data Fig. 5 in Okada et al.24). This might be explained by non-eQTL-SNPs with large effects, such as non-synonymous SNPs in genes PTPN22 (R620W) and TYK2 (P1104A), which exist in Europeans but are absent or exist to a lesser degree in Asian populations. Some eQTL-SNPs were estimated to have large effect size |*β*| > 0.05 (Fig. 2) in rheumatoid arthritis.

The SP-HMM can also provide posterior effect-size estimates of individual SNPs based on the estimated genetic architecture,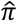 and 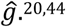^20,24^. To evaluate individual eQTL-SNPs, we used the estimated genetic architecture as the prior and listed the top SNPs with larger posterior means of effect size, |*β*| > 0.05 (Supplementary Data Set 1). As this list includes eQTLs such as RNASET2 and ADO, which have not been previously linked to rheumatoid arthritis^24^, this approach might be effective for identifying disease associated eQTL-SNPs. For the other diseases, enrichments of per-SNP variance due to the eQTLs in peripheral blood cells were also observed. Since eQTL-SNPs are associated with immune-related gene expression, these observations were consistent with the fact that coronary artery disease is a chronic inflammatory disorder and that genetic overlap between immune diseases and schizophrenia has been previously reported^45^. However, it should be noted that precise estimation of the eQTL effects in these diseases needs additional eQTL data covering all the tissues and cells related to the diseases.

Using DAF-stratified analysis for rheumatoid arthritis, we estimated more risk/protective derived alleles in low/high DAF (Fig. 3). Simple models based on a theory of population genetics for DAF46 (see Supplementary Fig. 20) could help interpret results from the DAF analysis, and thus provide another perspective on the difference among diseases (see Supplementary Note for details). Among such models, the ‘deleterious-risk and advantageous-protective mutation’ model with weak selection was best fitted for rheumatoid arthritis (Supplementary Fig. 21). Because most of the risk genes for rheumatoid arthritis are implicated in immune system regulation^24^, these low- and high-derived alleles would tend to skew individual’s immune function towards either deleterious or beneficial directions. Meanwhile, this skewing may result in breaking the balance between immunity and tolerance, leading to rheumatoid arthritis.

**Figure 3.**
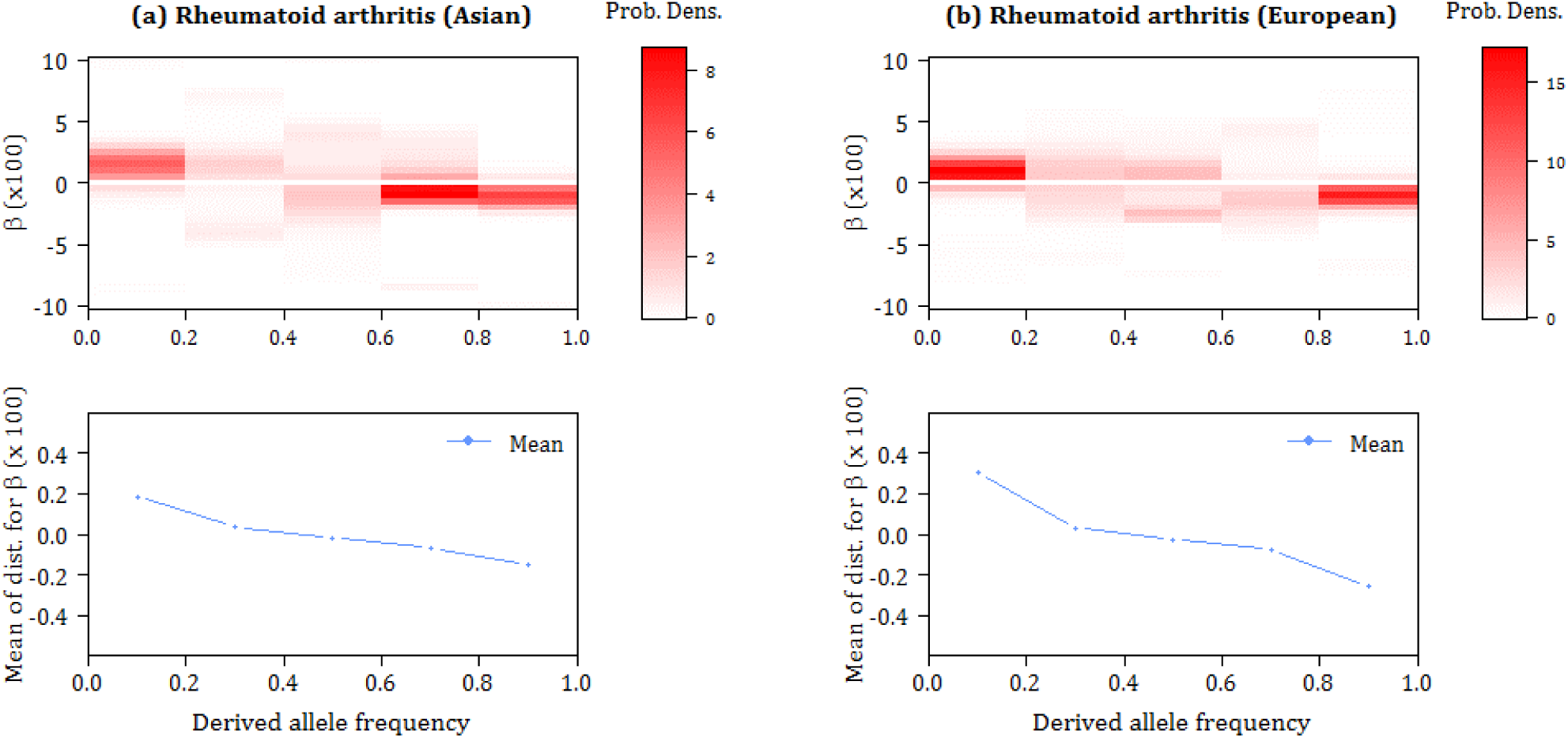
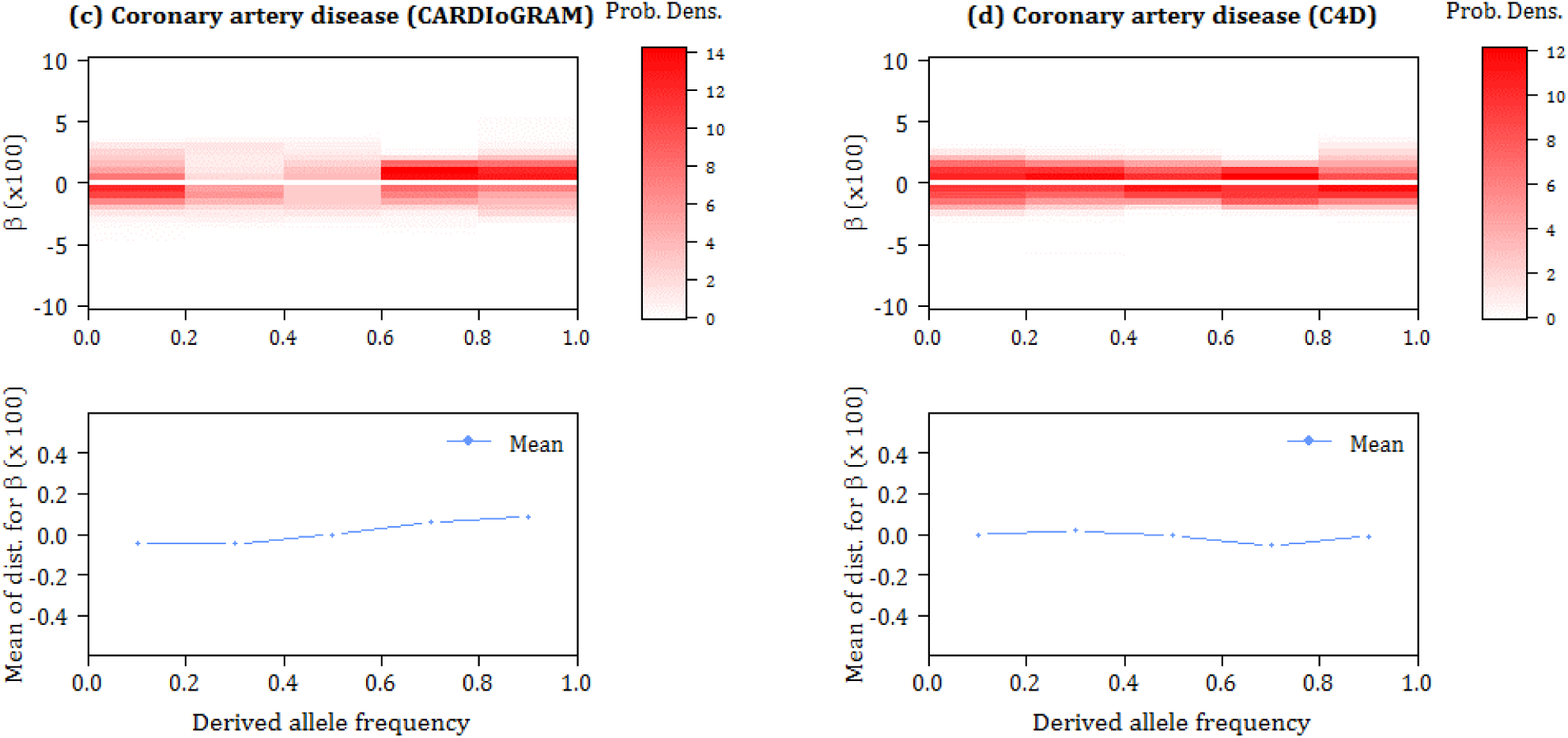
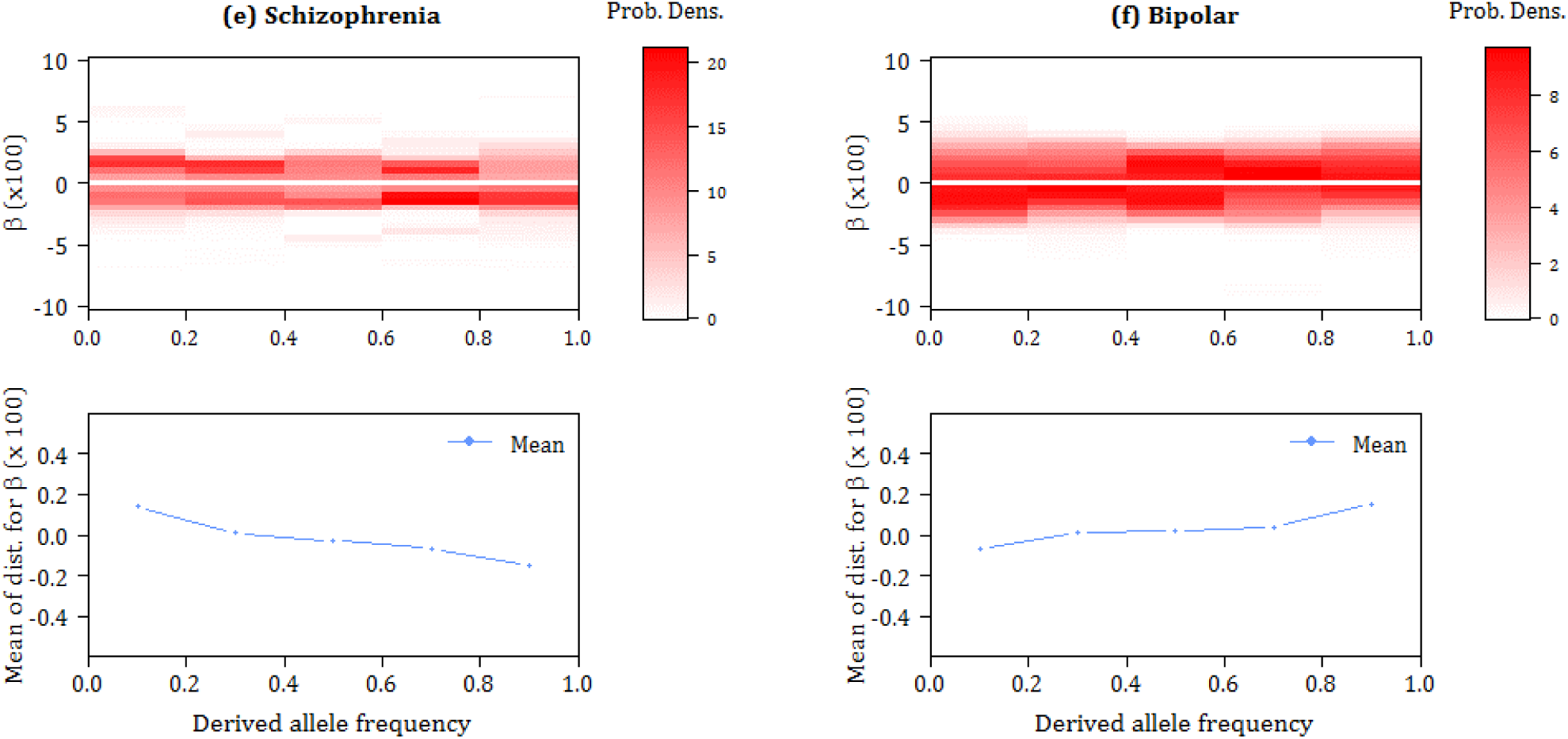
Estimated effect size distributions, 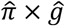, by derived allele frequency (DAF) bins. The upper panels (heatmap colors) for each GWAS results show 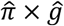. The lower panels show means of 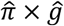.

Although some authors have reported that bipolar disorder and schizophrenia share a large amount of genetic factors^6,40^, we observed opposite tendencies in the genetic architecture for these diseases: risk (protective) and protective (risk) derived alleles were over-represented, especially at DAF ≤0.2 and DAF >0.8 for schizophrenia (bipolar disorder) (Fig. 3). This paradoxical result was consistent with a previous report that, among low minor allele frequency (1-5%) SNPs, the R/P ratio (ratio of the number of detected variants with risk in minor allele to those with protective effect) for schizophrenia was significantly larger than one, while for bipolar disorder it was less than one (see Table 1 in Chan et al. 35). Again, applying the same population genetics models, it was found that both the ‘deleterious-risk and advantageous-protective mutation’ and ‘deleterious-risk mutation’ models were better fitted for schizophrenia, whereas the ‘advantageous-risk and deleterious-protective mutation’ model was the best fitted for bipolar disorder (Supplementary Fig. 21). Recently, genetic correlations between creativity and both schizophrenia and bipolar disorder were reported, but they were much stronger for bipolar disorder^47,48^. Taken together with our estimation, these results might provide a clue for resolving the shared and specific genetic etiologies between the two genetically related diseases.

Lastly, the SP-HMM and empirical Bayes method, which can provide fine characterization of genetic architecture, can also contribute to accurate power analysis of GWAS^7,13^ and estimation of predictive capability of disease risk^15^. The SP-HMM can also be extended to multi-dimensional settings, e.g., for quantification of sex in genetic architecture for a disease, or (antagonistic) pleiotropic genetic architecture in multiple diseases. This kind of multi-dimensional analysis is novel and could provide new perspectives on multi-dimensional genetic effects, e.g., through a two-dimensional visualization of effect-size distributions for schizophrenia and bipolar diseases. Such analyses will be reported in future reports.

## URLs

- SP-HMM software, https://github.com/jonishino/SP-HMM
- HapMap 3, ftp://ftp.ncbi.nlm.nih.gov/hapmap/frequencies/2010-05_phaseIII;
- 1000 Genome, ftp://ftp-trace.ncbi.nih.gov/1000genomes/ftp/release/20130502;
- dbSNP (Build 141), http://www.ncbi.nlm.nih.gov/SNP;
- eQTL in blood, http://genenetwork.nl/bloodeqtlbrowser/2012-12-21-CisAssociationsP robeLevelFDR0.5.zip,
- rheumatoid arthritis summary statistics, http://plaza.umin.ac.jp/∼yokada/datasource/software.htm;
- schizophrenia and bipolar disorder summary statistics, www.med.unc.edu/pgc/downloads;
- coronary artery disease summary statistics, http://www.cardiogramplusc4d.org/.

## ONLINE METHODS

### Semi-parametric hierarchical mixture model (SP-HMM)

We defined the effect size, *βj*, for the *j*-th SNP of the total *m* SNPs as the genotype log-odds ratio under the additive allele dosage model. We considered the dosage of ‘derived mutant’ alleles. Namely, the genotypes *AA*, *Aa*, and *aa* in each SNP had dosages *xj* = 0, 1, and 2, respectively, where *a* was the derived and *A* was the ancestral allele. 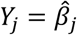 was an estimate of log-odds ratio for the *j*-th SNP (e.g., the standard maximum likelihood estimate). For *Y*_*j*_’s, we assumed a mixture structure with two components, null and non-null SNPs, in terms of association with disease susceptibility. To be specific,

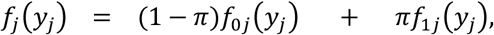

where *f*0*j* and *f*1*j* are the probability densities for null and non-null SNPs, respectively, and *π*is the prior probability of being non-null. For null SNPs, we specified 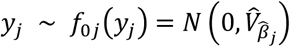 based on the asymptotic distribution of 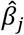, where 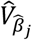 is an empirical variance estimate of 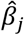 (e.g., the standard Wald-type variance estimate for 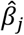). For non-null SNPs, we assumed the hierarchical structure: 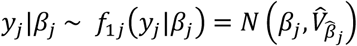 and *βj* ∼ *g*, where the prior effect-size distribution *g* was unspecified. In this model, the standard asymptotic normality was assumed for 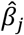 at the individual SNP level, while its true effect size *βj* followed a non-parametric prior distribution *g*, forming a semi-parametric hierarchical mixture model (SP-HMM)20,21. The assumption that *yj*’s are mutually independent would be reasonable for a set of LD-pruned SNPs.

### Empirical Bayes estimation

We estimated the priors, π and *g*, in the SP-HMM based on the data by applying an expectation– maximization (EM) algorithm, called the smoothing-and-roughening algorithm22, to incorporate the non-parametric prior distribution *g*20,21. The non-parametric estimate of *g* was supported by fixed discrete mass points ***p*** = (*p*1*, p*2*, …, p*_*B*_) at a series of nonzero points ***b*** = (*b*1*, b*2*, …, b*_*B*_) (*b*1 < *b*2 <…< *b*_*B*_). We specified a wide range for the mass points, such as *b*1 = –0.3 to *b*_*B*_ = 0.3 (0.74 to 1.35 in odds ratio), to support the effect-size distributions in many complex diseases. We set the number grid points as 120, such that ***b*** = (–0.300, –0.295, …, –0.005, 0.005, …, 0.295, 0.300). The initial value of π, π*init*, and the initial distribution of *g*, *ginit*, were determined sequentially. Setting *g* to be uniformly distributed (i.e., *p*_*i*_=1/*B* for all *i*), the EM procedures for candidate initial values, π = 0.1, 0.2, …, or 0.9, were ran 200 times and the value of estimated π with maximum likelihood was selected as π*init*. Then setting *g* to be uniformly distributed again, we got *ginit* by the EM procedure with fixed π = π_*init*_(the EM iterations were stopped when the relative change of π in one iteration was small < 0.005 % or until 200 iterations). Setting *g* = *g*_*init*_ and π = π_*init*_, the final (main EM algorithm was stopped after at least 2000 iterations when the relative changes in the estimate of *π* in one iteration were small (< 0.005 %) or iterations reached 2000 times. We applied a parametric bootstrap method based on the estimated SP-HMM to estimate standard errors of the estimate for π.

### Liability-scale variance explained by SNPs

As shown by So et al 49, the log odds ratio, *β*_*j*_, together with the allele frequency and the disease prevalence, were transformed to the variance explained by the *j*-th SNP, denoted as *v*_*j*_, in the liability threshold model. In the liability threshold model, we assumed that an underlying liability to disease follows a normal distribution and individuals that exceeded a threshold of liability, *T*, were affected with the disease. Individuals with the genotypes of *AA*, *Aa*, and *aa* at the *j*-th locus had liability distributions with different means, but the same residual variance. We let *p*_*j*_ be the derived allele frequency and *h*_*j*_,*x*_*j*_ be the frequency of genotype *x*_*j*_ (*x*_*j*_ = 0, 1, 2) in the general population. Assuming the Hardy-Weinberg equilibrium in the population, the genotype frequencies are given by *h*_*j*,0_ = (1 - *pj*)^2^, *h*_*j*,1_ = 2*pj* (1 - *pj*), and *h*_*j*,2_ = *pj* 2. Using the overall mean liability, *μ*_*all*_, and the mean liabilities of genotype *xj*, *μ*_*j*_,*x*_*j*_, the variance explained by *j*-th SNP is given by

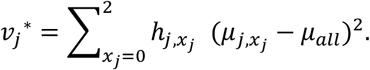

For evaluating *μ*_*j*,*xj*_, we used the penetrance of genotype *x*_*j*_, denoted by *φ*_*j,x,j*_ = 1/(1 + *e*^−*α*_*j*_−*β*_*j*_*x*_*j*_^ under the additive allele dosage model, where α*j* was determined under the constraint involving the disease prevalence *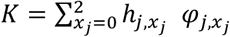*. Assuming that the residual variance of each genotype was 1, the mean liability of each genotype was given by

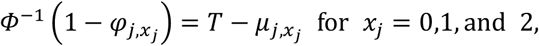

from which we obtained values of *μ*_*j,x,j*_, where *Φ*was the cumulative distribution function of the standard normal distribution. Of note, one of the mean liabilities of genotypes can be set as an arbitrary value, as it does not affect the variance estimate. Finally, *v*_*j*_ was obtained by *ν*_*j*_ = *ν*_*j*_^*^/(1 + *ν*_*j*_^*^). This corresponded to the variance under the standard liability threshold model with the unit total variance of liability, as is assumed in heritability estimation^50,4^

We estimated the distribution of *v*_*j*_ for non-null effects using the estimated effect-size distribution 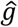, together with using allele frequencies and the prevalences. The allele frequencies were retrieved from the 1000 Genome phase III51 and the same prevalences as previously assumed in estimating SNP heritability were used8,9. Then, the point estimate of *v*_*j*_, 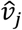, was gained as the product of the estimate 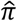 and the mean of the estimated distribution of *v*_*j*_ for non-null effects. The total liability-scale variance, *V*, explained by the pruned estimated as a simple sum of 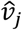 over all SNPs in the sets.

### GWAS data analysis

The six sets of GWAS summary statistics that we used were available online (see URLs). The characteristics of individual GWASs are shown in Supplementary Tables 1 and 2. For rheumatoid arthritis, the MHC region (chromosome 6, 25 – 35 Mb) was removed. The derived/ancestral states of alleles were determined by using the dbSNP.

We used two kinds of pruned SNP sets, P-value-based and random-pruned sets, in the non-stratified SP-HMM analysis (Table 1 and Fig. 1). To gain the P-value-based pruned set for a GWAS, we began by selecting the most strongly associated SNP, i.e., the SNP with the lowest P value, in a reference GWAS as a SNP of the pruned set, and all other SNPs in LD (*r*^2^ > 0.1) with the selected SNP were removed. The process was repeated until no SNPs remained. LD information was retrieved from the HapMap data base (HapMap phases I+II+III, release 27)52. In selecting SNPs with strong associations for Asian rheumatoid arthritis GWAS, European rheumatoid arthritis GWAS data were used as a reference for association, and vice versa. For coronary artery disease, the data of two GWAS, CARDIoGRAM and C4D, were used reciprocally. For the two genetically correlated diseases, schizophrenia and bipolar disease, the data of two GWAS for the two diseases were used reciprocally. For the random-pruned sets, we included SNPs randomly, irrespective of degrees of association, i.e., P values in the reference GWAS data, such that no SNPs in the set were in *r*^2^ > 0.1.

For stratified analysis by eQTL/non-eQTL-SNPs, we defined ‘eQTL SNP’ as cis-eQTL SNPs detected with false discovery rate < 0.5 using peripheral blood samples (Westra et al., 2013). In the eQTL/non-eQTL-SNPs set analyzed, all the eQTL and non-eQTL SNPs were selected to be nearly independent of one another (*r*^2^ ≤0.1). In this data set, eQTL SNPs showing stronger associations (i.e., lower *P* values) with gene expressions were preferentially included, and LD pruning was conducted as in the P-value-based pruned sets. Non-eQTL SNPs were randomly selected.

In DAF-stratified analysis, the allele frequencies of SNPs were determined by the 1000 Genome phase III data51. For each DAF bin, we used 100,000 SNPs randomly selected from GWAS SNPs regardless of LD. This was because estimates of SP-HMM were unstable due to small number of SNPs (e.g., a few thousand SNPs) when LD pruned sets were used. Note that, in C4D GWAS, the numbers of SNPs used in 0.4 < DAF ≤0.6, 0.6 < DAF ≤0.8, and 0.8 < DAF were 94506, 70170, and 49116, respectively, since the SNPs of C4D GWAS was limited (Supplementary Table 2). The obtained results (i.e., estimates of *π* and *g*) using the pruned sets (data not shown) were close to those sampled regardless of LD, and both results had the same trends over DAF bins.

For selecting high quality SNPs and LD information in the above section, HapMap data of Japanese individuals in Tokyo (JPT) and European-ancestry individuals from Utah (CEU) were used for Asian rheumatoid arthritis GWAS data and the other GWAS data, respectively. Similarly, for information of allele frequencies, East Asian and European 1000 Genome Project data were used for Asian rheumatoid arthritis GWAS data and the other GWAS data, respectively.

## Acknowledgments

This research was supported by JST CREST Grant Number JPMJCR1412, Japan and a Grant-in-Aid for Scientific Research (16H06299) from the Ministry of Education, Culture, Sports, Science and Technology of Japan.

## Author Contributions

J.N. developed the methods, performed the analyses, and wrote the manuscript. Y.K. provided essential ideas and interpretations for the study direction and results. D.S. and T.M. contributed to the data acquisition and the analyses. H.M. provided the initial version of script for SP-HMM analysis. Y.K., M.K., H.O., K. A. B., and T.T. improved the manuscript. T.T. directed and supervised the study. S.M. conceived the study idea, developed the methods, wrote the manuscript, and, directed the study. All authors contributed the final manuscript.

## Competing Financial Interests

The authors declare no competing financial interests.

